# Shoot to root transport of the HY5 transcription factor drives root developmental plasticity under neighbour shade

**DOI:** 10.64898/2026.10.18.759902

**Authors:** Kyra van der Velde, Ahmad Ali, Valérie Hoogers, Franziska Stamm, Alena Patnaik, Omar Heliel, Douwe Rietveld, Tuğba Akyuz, Katarzyna Retzer, Jan Petrašek, Ronald Pierik, Kasper van Gelderen

## Abstract

Plants perceive competitors through the reflection of Far-Red light from neighbouring leaves. A decrease in the Red: Far-red light ratio (R:FR) is perceived by phytochrome photoreceptors and triggers elongation responses aboveground, allowing plants to anticipate future competition for light; their primary energy source. However, low R:FR perception by the shoot also prompts an alteration of root development, possibly to conserve resources. An important regulator of this response is the bZIP transcription factor ELONGATED HYPOCOTYL 5 (HY5), which can be transported from the shoot to the root, where it suppresses auxin signaling and lateral root outgrowth. However, it remains unknown whether HY5 transport is the direct link between shoot-perceived low R:FR and a decrease in lateral root outgrowth. We therefore sought to uncover the mechanics and speed of HY5 transport. To this end, we constructed shoot-specific inducible lines to perform live-imaging of HY5 transport using a vertical confocal microscope. This allowed us to follow the transport and unloading of HY5 from the phloem into the protophloem pericycle, where it becomes nuclearly localized, in real time over a 24-hour period. To mimic plant competition, we applied supplemental FR light and observed that this increased the HY5 protein levels in the shoot and subsequent HY5 protein transport from shoot-to-root. Finally, we were able to show that shoot-expressed HY5 is necessary and sufficient to induce the root response to shoot perceived Far-Red light. This work showcases in detail how a small general transcription factor functions as a long-distance signal between shoot and root and integrates environmental signalling over long distances in the plant.

## Introduction

Plants compete for limiting resources while growing in dense vegetation. While water and nutrients are primarily taken up by the roots, aboveground tissues compete for light. In high-density vegetation, plants compete for photosynthetically active radiation, including red light (R) while reflecting far red light (FR), thereby reducing the R:FR in the canopy. This R:FR ratio is sensed by phytochrome photoreceptors. Phytochromes exist in two interconvertible forms: the inactive red-light-absorbing Pr form and the active far-red-light-absorbing Pfr form. In standard sunlight R:FR conditions, phytochrome B (phyB) is predominantly present in the active Pfr form, which represses Phytochrome Interacting Factors (PIFs) by promoting their phosphorylation and degradation. Under low R:FR conditions, phyB photoconversion to the inactive Pr form releases suppression of PIFs (Franklin & Whitelam, 2005; Ni *et al*, 2013; Franklin & Quail, 2010), allowing them to accumulate and activate downstream shade avoidance responses, including upward leaf movement, stem elongation, and early flowering (Ballaré & Pierik, 2017).

While photosynthesis and light competition occur in aerial tissues, low R:FR ratios also affect root development. Low R:FR reduces lateral root development, by decreasing the emergence of lateral root primordia (LRP) from the primary root (van Gelderen et al. 2018). This effect requires a mobile signal, as the response depends on phytochrome action in the shoot, yet roots typically grow in dark soil and light perception is therefore restricted to aerial tissues. Shoot-to-root mobile signals can travel through the phloem and can include a diversity of molecules, including phytohormones, proteins, peptides and RNA molecules. In the case of low R:FR, specific mobile signals identified in the past are the hormone gibberellin (van Gelderen et al. 2025), and the bZIP transcription factor ELONGATED HYPOCOTYL 5 (HY5) (Chen et al. 2016; Sakuraba et al. 2018; van Gelderen et al. 2018), and both are proposed to move from shoot to root through the phloem. Phloem loading involves both active transporters and via plasmodesmata (Zhang and Turgeon 2018). Small molecules move freely, while larger molecules require passive diffusion through enlarged plasmodesmata or active loading mechanisms. Phloem loading is therefore limited by the size exclusion limit of plasmodesmata. In Cucumber phloem loading, much larger exclusion limits of plasmodesmata have been observed than in *Arabidopsis*, increasing to 20 to 40kDa (Balachandran et al. 1997). Phloem unloading at the root tip is thought to occur by a combination of bulk flow and passive diffusion (Ross-Elliott et al. 2017). At the phloem end near the root tip, proteins and solutes exit the protophloem into the phloem-pole pericycle (PPP). The size exclusion limit of plasmodesmata at these sites is thought to determine which proteins can exit the phloem (Ross-Elliott et al. 2017). While small molecules are unloaded from the phloem without restriction, larger proteins undergo batch unloading exclusively into the PPP (Paultre et al. 2016; Ross-Elliott et al. 2017). Between the phloem and the PPP, larger plasmodesmata have been observed with high exclusion limits up to and exceeding 120kDa (Ross-Elliott et al. 2017).

The bZIP transcription factor HY5 can suppress lateral root emergence under low R:FR conditions. In the root, HY5 acts as a transcriptional repressor of lateral root development through direct repression of the expression of Auxin Response Factor 19 (ARF19), and auxin transporters PIN-FORMED3 (PIN3) and LIKE AUX1 3 (LAX3) (van Gelderen et al. 2018), thereby affecting local auxin signalling and transport. Beyond lateral root development, HY5 shoot-to-root transport has also been implicated in regulating nitrate and phosphate uptake from the soil, integrating light quality signals with nutrient acquisition (Chen et al. 2016; Sakuraba et al. 2018; van Gelderen et al. 2021). HY5 transport likely happens through the phloem (Chen et al. 2016), but despite growing evidence that HY5 functions as a shoot-to-root mobile signal, it is unknown how HY5 is transported. HY5 might enter the phloem in the leaves or cotyledons and be transported toward the root tip. The rates of HY5 transport from shoot-to-root and the functional significance of this transport for lateral root development remain to be resolved. Addressing these knowledge gaps requires tools that allow HY5 transport to be visualised in real time.

Therefore, we generated HY5 shoot-specific inducible lines to follow HY5 transport in real time by using a vertical confocal microscope setup to monitor HY5 phloem unloading at the root tip. We found that HY5 unloading occurs at the end of the protophloem within hours after induction in the shoot, and that the amount of unloaded HY5 is enhanced under low R:FR. Low R:FR also stabilizes HY5 in the shoot and shoot expression of HY5 is sufficient to recapitulate the low R:FR lateral root phenotype. Finally, although shoot-derived HY5 represents only a small fraction of the total HY5 pool in LRP, we show that shoot-derived HY5 is responsible for the elevated HY5 accumulation in LRP under low R:FR, despite being present at only low levels in these tissues.

## Results

### HY5 is unloaded into protophloem pericycle

To confirm shoot-to-root HY5 translocation (Chen et al. 2016), we grafted a *hy5-2 35S:HY5-GFP*scion onto a Col-0 rootstock. We observed a clear nuclear GFP fluorescence signal in two cell files, the pericycle, near the phloem unloading zone (Figure 1A; arrowheads), together with a relatively low signal in the cortex. This signal was reproducible across grafts, whereas self-grafted Col-0 rootstocks showed no comparable nuclear signal (Figure S1). Grafting a *hy5-2 pHY5:HY5-GFP* scion onto a Col-0 rootstock resulted in a weaker but similarly visible signal throughout the root tip (Figure 1B), confirming that HY5 produced in the shoot under its native promoter, reaches the root.

**Figure 1.**
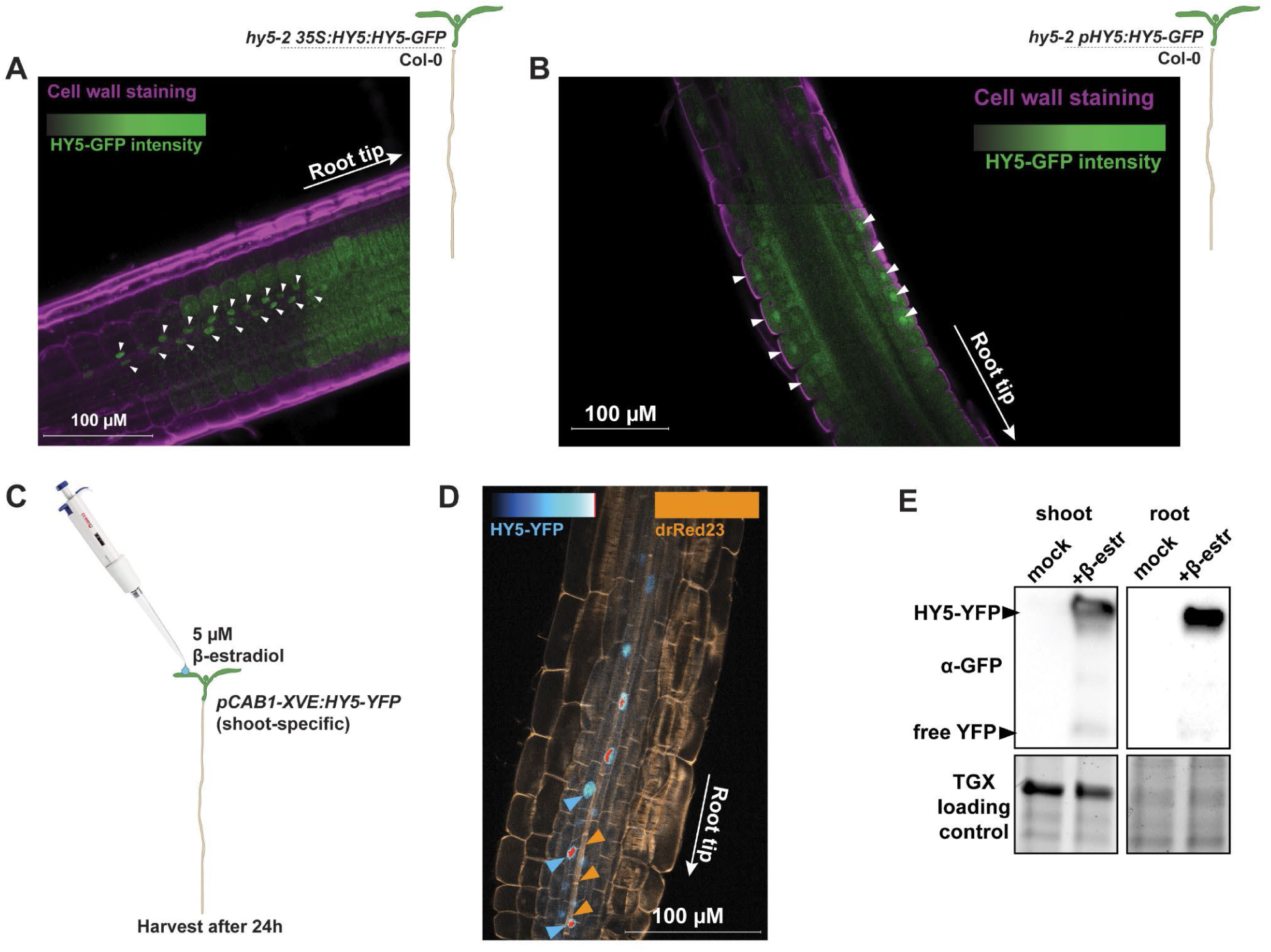
HY5 is transported from the shoot and unloaded close to the root tip. (A) Elongation zone of grafted seedlings with *hy5-2 35S:HY5-GFP* scion and Col-0 wild type rootstock. (B) Elongation zone of grafted seedling with *hy5-2 pHY5:HY5-YFP* scion and Col-0 wildtype rootstock. (C) Graphic showing induction of shoot-inducible *pCAB1-XVE:HY5-YFP* with 5 μM ꞵ-estradiol on the shoot, 24h before harvest for Western blotting. (D) image of fixed sample of *pCAB1-XVE:HY5-YFP,* induced by the method of (C), cell walls were stained with direct-red 23 (drRed23), orange arrowheads point out the newly-formed phloem strand, which stains brighter with drRed23, while the blue arrowheads indicate the presence of HY5-YFP signal on top of the phloem. (E) Western blot of Col-0 *pCAB1-XVE:HY5-YFP* lines, treated with mock or ꞵ-estradiol and WL, with shoot and root samples probed with anti-YFP antibody (Representative experiment of three independent replicates).

While grafting confirms shoot-to-root movement, it does affect seedling development and the graft-junction potentially affects transport over the junction. To follow HY5 transport in real time in intact seedlings, we generated several HY5-YFP inducible lines driven by the shoot-specific *pCAB1* promoter combined with the XVE cassette ꞵ-estradiol induction system (Siligato et al. 2016). Because *pCAB1* drives expression specifically in the shoot, any observed HY5-YFP signal in the root can be attributed to transport rather than expression in the root. We applied a 0.1% agarose droplet containing 5μM ꞵ-estradiol to the shoot of these seedlings and sampled shoot and root tissues for microscopy and western blot analysis (Figure 1C). Examination of the microscopy signal in combination with a cell wall stain showed in more detail that the signal is localized in two-cell files above the newly formed phloem strands (Figure 1D). This signal is the protophloem pericycle (PPP), from which the phloem unloads its cargo (Ross-Elliott et al. 2017). The protophloem pericycle is the first cell layer around which the phloem is differentiated, representing the effective end of phloem transport. Through western blot analysis we observed that intact HY5-YFP was detected in the root of induced seedlings but was absent in mock-treated controls (Figure 1E), confirming that HY5-YFP is transported from shoot to root in the inducible system.

### HY5 unloads in distinct pulses, starting after 6 hours of shoot-induction

We next imaged the shoot-specific, inducible *HY5-YFP* lines with a vertically mounted confocal microscope (von Wangenheim et al. 2017), to track HY5-YFP arrival in the root over time. Seedlings were induced with estradiol on the shoot at noon, 6 hours prior to imaging, and then imaged for 15 hours with 20 minute intervals. HY5 signals were visible from 6 hours after induction, and onwards (Figure 2A). In general, the HY5-YFP signal was preceded by a diffuse signal in the phloem strand, followed by the appearance of nuclear signal in the PPP, in sudden, bursts, similar to free GFP unloading from the phloem (Figure 2A, supplementary movie 1) (Ross-Elliott et al. 2017). This nuclear signal persisted throughout the experiment (Figure 2A, green arrowheads). The HY5-YFP signal was mostly observed in the two PPP cell files directly connected to the phloem (Figure 2B) (Ross-Elliott et al. 2017). During unloading, nuclear signal appeared within 30 minutes of the onset of an unloading event (Figure 2C, supplementary movies 1&2).The HY5 signal then persisted in these cells and was not observed to spread further. Mock treatment with DMSO instead of beta-estradiol yielded no clear HY5-YFP signal, nor did direct estradiol treatment of the root (Figure S2, supplementary movie 3). An inducible *HY5-YFP* construct in the *hy5-215* mutant background produced a similar HY5 transport pattern (Figure S3, supplementary movie 2).

**Figure 2.**
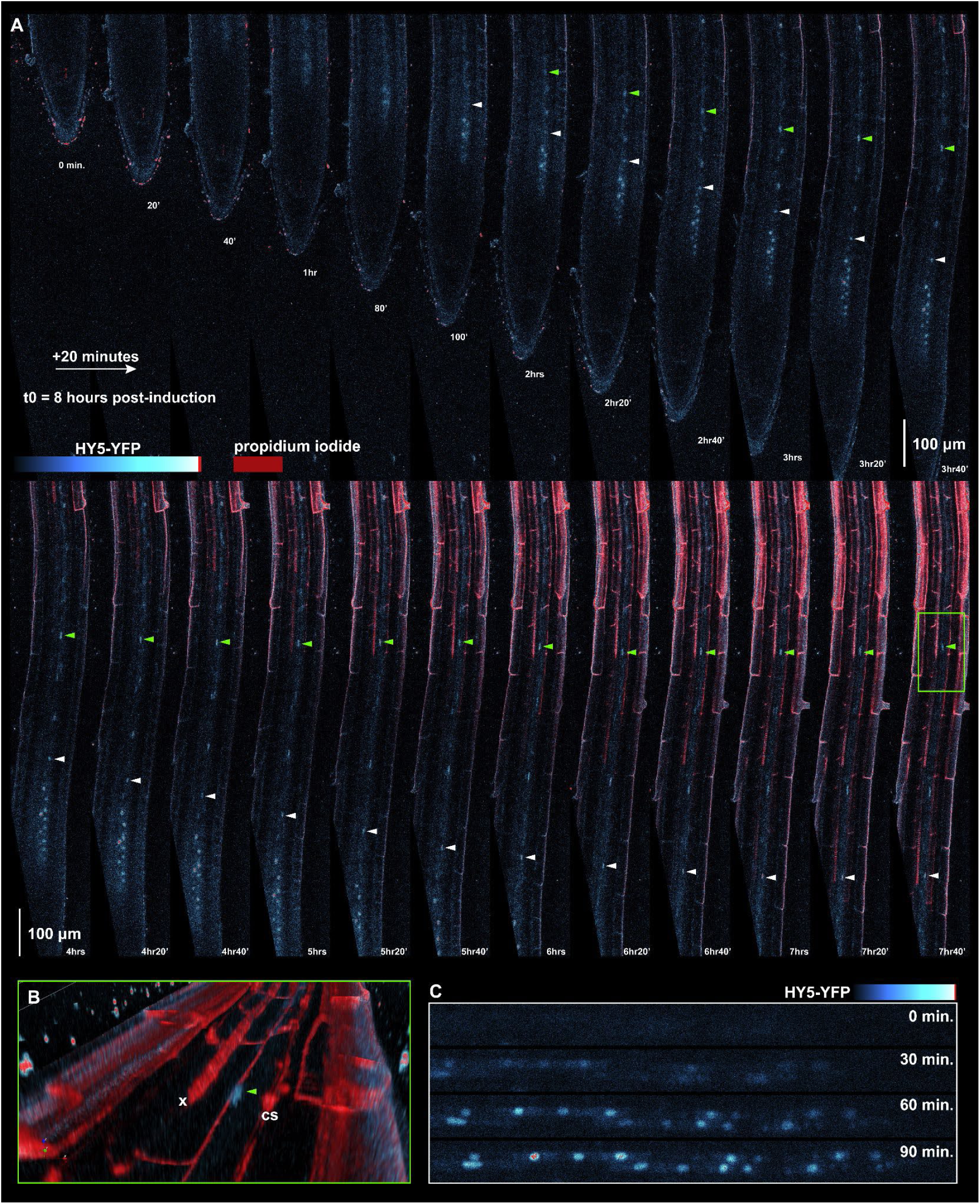
HY5 is unloaded from the phloem into the protophloem pericycle in distinct pulses. (A) Representative image of confocal time-lapse live imaging experiment using Col-0 *pCAB1-XVE:HY5-YFP* inducible lines. HY5-YFP was induced in the shoot, 6 hours prior to the start of imaging (number of replicates, 14). Seedling root tips were imaged using a 90 degree tilted confocal setup (zeiss lsm 880), and followed for 15 hours. Green and white arrowheads show HY5 nuclear signals persisting into the differentiation zone and elongation zone. For original video, see supplemental movie 1, and *hy5-215-pCAB1-XVE:HY5-YFP* background see supplemental movie 2. (B) 3D representation of root from (A) showing localization of unloaded HY5 (green arrowhead) (x = xylem, cs = Casparian strip). (C) Detail showing the appearance of HY5-YFP signal in the nuclei of the root tip PPP.

### FR light perception increases shoot-HY5 and its shoot-to-root transport

In previous work, we uncovered the role of *HY5* in regulating the root response to shoot-perceived far-red light (van Gelderen et al. 2018, 2021, 2025). Therefore, using *hy5-2 hyh pHY5:HY5-YFP* lines we first investigated the effect of low R:FR on HY5 in the shoot. Phytochromes are known to inhibit the action of COP1, the E3 ligase that binds SPA proteins and mediates HY5 degradation (Lu et al. 2015; Sheerin et al. 2015), while there is an indication that FR light can stabilize HY5 (Pacin et al. 2016). We used supplemental far-red (white light + far-red) to achieve low R:FR conditions, henceforth called WL+FR. We treated 3-day-old seedlings with WL, WL + far-red (WL+FR) for 24 hrs and WL+FR for 48 hrs, and harvested the seedlings at 4 and 5 days old. Quantification of HY5-YFP in confocal images showed that both the 24 and 48 hr WL+FR treatments led to an increase in HY5-YFP in the leaf primordia, shoot apical meristem and the hypocotyl vasculature (Figure 3).

**Figure 3:**
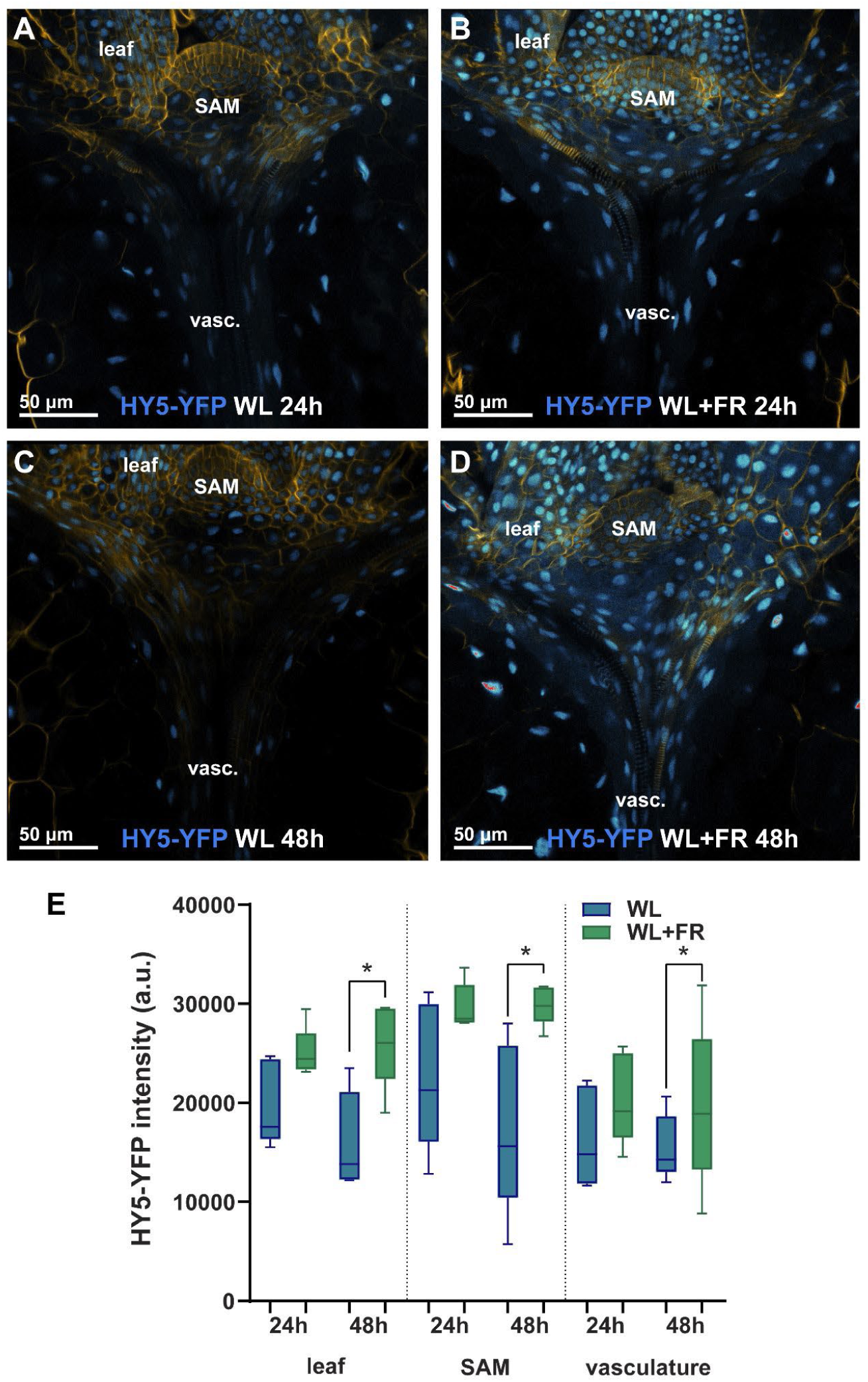
Supplemental FR treatment increases HY5-YFP accumulation in young shoot tissues and vasculature. (A–D) Representative confocal images of *hy5-2 hyh pHY5:HY5-YFP* seedlings showing HY5-YFP signal in leaf primordia, shoot apical meristem (SAM), and shoot vasculature. Seedlings were treated with WL (A, C) or WL+FR (B, D) for 24 h (A, B) or 48 h (C, D), and fixed at 4 or 5 days old, respectively. Samples were cleared with ClearSee and cell walls stained with Direct Red 23 (shown in orange-yellow). (E) Average pixel intensityof HY5-YFP in leaf primordia, SAM, and vasculature across conditions (n=10).

To test whether this increase of HY5 in the shoot translates into increased delivery to the roots, we repeated the live imaging experiments described for Figure 2 but extended the white light pre-treatment (120 μmol m^-2^ s^-1^) with a supplemental FR pre-treatment (WL+FR 120 μmol m^-2^ s^-1^, R:FR 0.2). Similar to the experiments in Figure 2, mock treatment with DMSO did not result in HY5 signal (Figure 4A, Movie S3), whereas induction with estradiol did. Because the timing of each HY5 transport event varied, we quantified the time window with the strongest signal in each replicate. This yielded at least 19 individual roots with HY5 transport per treatment. Across the entire experiment, roughly 85% of roots had a HY5 transport event (Figure S4). WL+FR pretreatment resulted in a significantly higher intensity and number of HY5 transport events than observed in WL (Figure 4B,C, S4, Movie S4,5).

**Figure 4:**
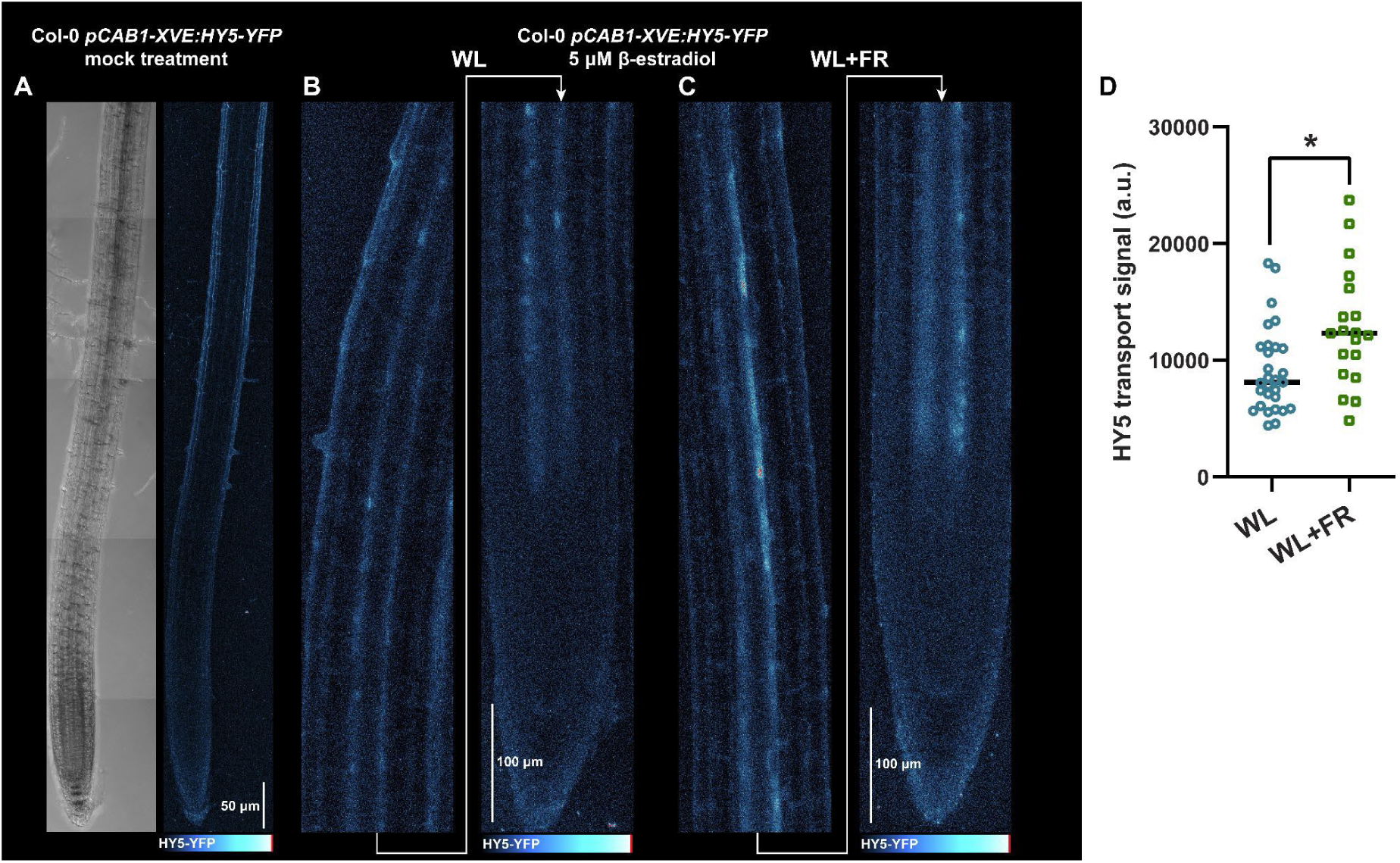
Supplemental far red light increases HY5 transport and unloading. (A) Representative stills of confocal live imaging series of mock and ꞵ-estradiol treated in WL and WL+FR. (B) Quantification of HY5 signal intensity, minus background signal from the dataset from (A) (n=36 WL, 25 WL+FR).

### HY5 protein, not mRNA, is transported to the root

Confocal imaging revealed the HY5 spatial distribution in shoot tissues and HY5 transport to root tissues in WL and WL+FR. To confirm HY5-YFP induction and transport at the protein level, we performed Western blots on shoot and root samples of the pCAB1 inducible lines in Col-0 and *hy5-215* background. Estradiol-induced expression of HY5-YFP in the shoot was confirmed by a strong band in shoot samples under WL, and WL+FR (Figure 5A,B). Importantly, HY5-YFP was also detected in the root samples, with enhanced accumulation under WL+FR compared to WL (Figure 5A,B), consistent with increased transport and unloading shown in Figure 4C. In the *hy5-215* mutant background we observed similar induction and transport as in Col-0 background, and here we observed increased HY5 levels in WL+FR after induction, as well as clear shoot to root transport in WL+FR, and not in WL (Figure 5C). This further indicates that HY5 transport relies on stabilization of the protein, and not on *HY5* transcription, and that this stabilization promotes the transport (Figure 5C).

**Figure 5:**
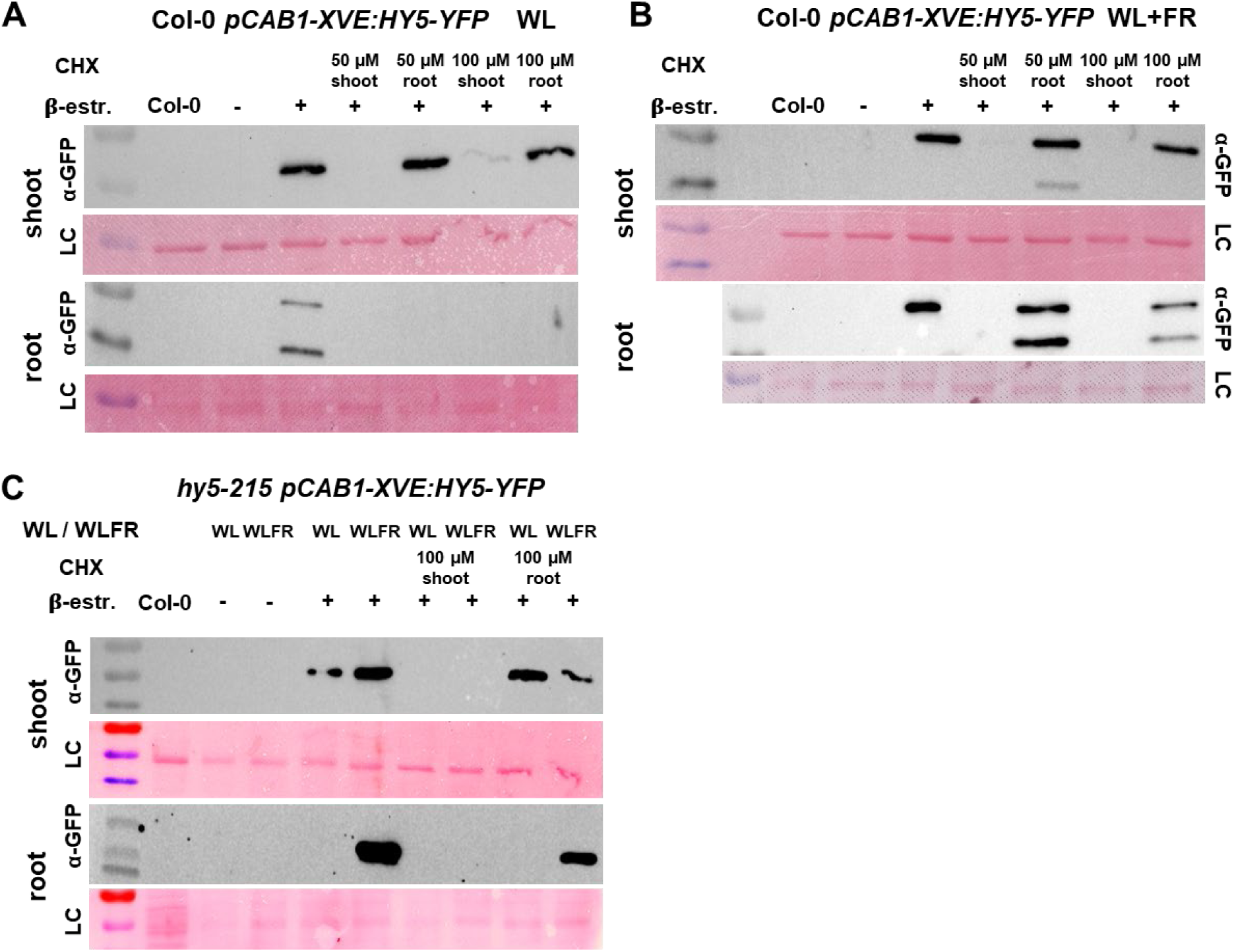
HY5 transport is not dependent on protein translation in the root. Western blot on seedling roots or shoots from HY5 shoot-inducible lines. (A and B) Col-0 *pCAB1-XVE:HY5-YFP* lines were induced with 5 µM beta-estradiol on the shoot, and additionally the translation inhibitor cycloheximide (CHX) was applied on the shoot or root. After 24 hours, seedlings treated with either WL (A) or WL+FR (B) were cut at the shoot/root junction and frozen in liquid N. Western blot membranes were probed with anti-GFP and stained with Ponceau afterwards as loading control (LC). The upper detected bands are around 55kDa, while the lower bands are around 40 kDa. (C) Similar experiment as (A and B), but with a *hy5-215 pCAB1-XVE:HY5-YFP* inducible line (Blots are representative of three independent replicates).

To verify that the observed HY5 signal indeed originates from protein transport and not from transported *HY5* mRNA getting translated in the roots, we treated the shoot or the root tip with the translation inhibitor cycloheximide (CHX) while simultaneously inducing *HY5-YFP* with beta-estradiol in both the Col-0 and *hy5-215* background (Figure 5, right side of blots). Application of either 50 or 100 µM CHX to the shoot was sufficient to block induction of HY5, thereby preventing subsequent transport to the root (Figure 5A,B,C). In contrast, application of the same CHX concentrations to the root did not block HY5 transport under WL+FR (Figure 5B,C). Root-applied 100 µM CHX did result in fainter bands in the root compared to 50 µM CHX. CHX in the roots led to a second GFP band to emerge. The weight difference between the two bands suggests that this could be due to the effect of ubiquitination, induced by phosphorylation or other post-translational modifications that are common for HY5 (Hardtke et al., 2000, Hoecker et al., 2017). In a parallel experiment, seedlings were treated in exactly the same manner, but were fixed with and cleared for confocal microscopy. We observed HY5 signal at the phloem poles in a similar manner as Figure 2, and this signal was absent or strongly reduced when translation was blocked at the shoot, but not the root (Figure S5). Taken together, these results indicate that mRNA translation in the root is not required to observe shoot-induced HY5 to accumulate in the roots, confirming that it is very likely the protein itself that is moving from shoot-to-root.

### Shoot-expressed *HY5* rescues WL+FR-induced root growth reduction

We next sought to establish a functional link between three observations: the WL+FR-induced reduction of lateral root development, which is regulated by HY5 (van Gelderen *et al*, 2018), increased HY5 protein in shoot tissues by WL+FR, and the increased rootward HY5 translocation in WL+FR. To this end, we used constitutively shoot-expressed HY5 lines (*pCAB1:HY5*), within wildtype and *hy5-215* mutant backgrounds to assess shoot and root phenotypes in WL and WL+FR. The *pCAB1:HY5* line had a regular hypocotyl length in both WL and WL+FR, resembling the *HY5* overexpression line (Figure 6A)(van Gelderen *et al*, 2018). *hy5-215* had an elongated hypocotyl under WL but still responded to WL+FR and the *pCAB1:HY5* construct in *hy5-215* partially complemented it (Figure 6A). Lateral root density in Col-0 was reduced under WL+FR, and this reduction was absent in *hy5-215* (Figure 6B). In the Col-0 background, the *pCAB1:HY5* construct produced a *HY5* overexpression phenotype, however it was able to reconstitute the wild type phenotype in the *hy5-215* background, showing the importance of shoot-expressed *HY5* for regulating lateral root development (Figure 6).

**Figure 6:**
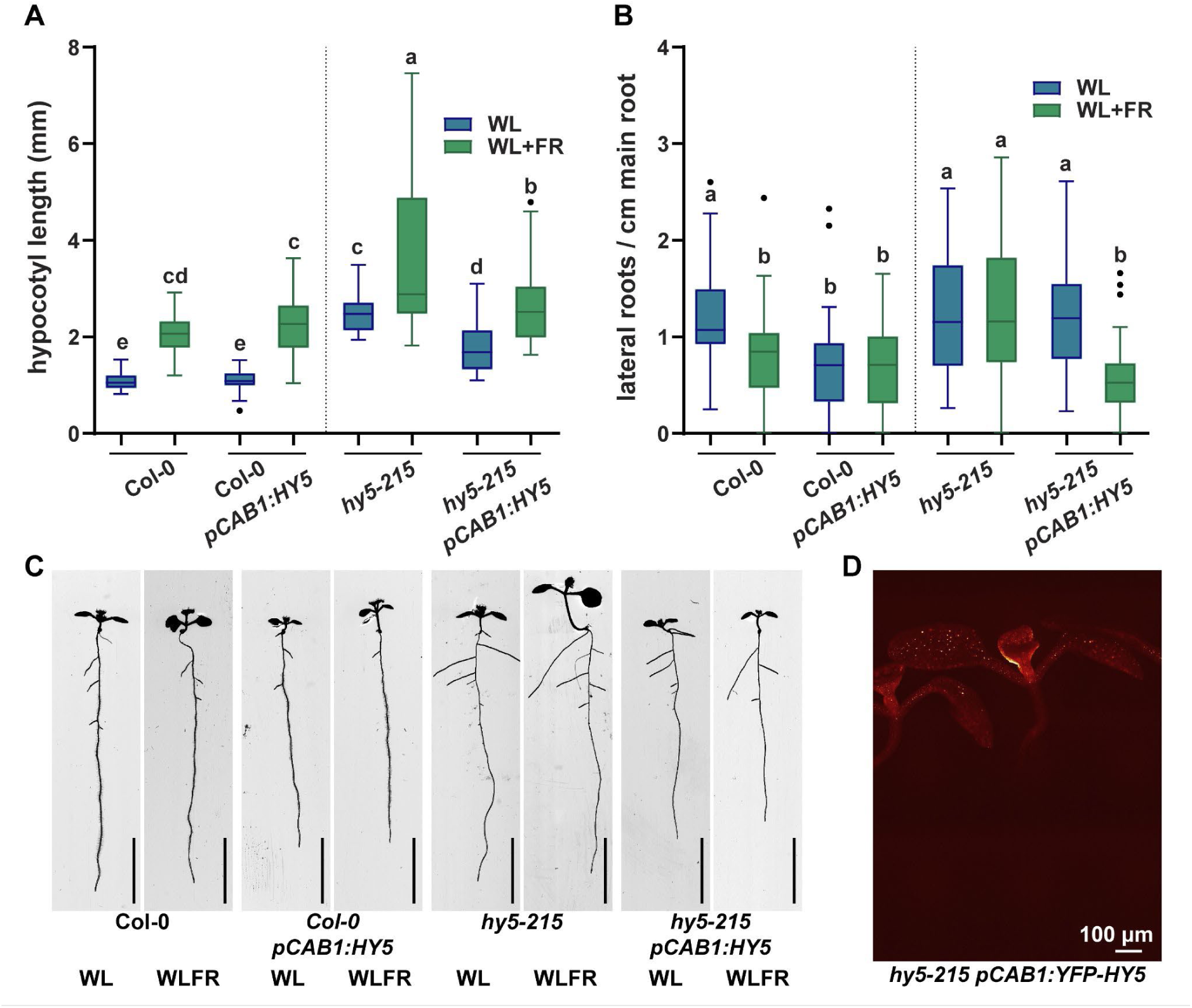
Shoot-expressed HY5 rescues *hy5* mutant loss of WL+FR phenotype. (A) Hypocotyl length and (B) Lateral root density of lines with a *pCAB1-*driven *HY5-YFP* construct in Col-0 or *hy5-215* mutant background. Seedlings were grown in either WL (130 umol/m2/s), or WL+FR (R:FR ratio 0.2). Statistical differences are indicated by the letters above the boxplots (two-way ANOVA p<0,05, n= 20<40)). (C) Representative images of experiment in (A,B). (D) Fluorescence stereoscope image of *hy5-215 pCAB1:YFP-HY5*.

### HY5 accumulation in lateral root primordia requires shoot-HY5

The *pCAB1:HY5* phenotypes in Figure 6 establish that shoot-expressed HY5 is sufficient to restore the lateral root density response to WL+FR. HY5 accumulation in lateral root primordia has previously been shown to drive the reduction in lateral root outgrowth under shoot-perceived WL+FR (van Gelderen et al. 2018). To test whether HY5 accumulation is dependent on shoot-borne HY5, we grafted Col-0 and *hy5hyh* shoot scions onto *hy5-2 pHY5:HY5-GFP* rootstocks. This allowed us to follow root-expressed HY5-GFP in the lateral root primordia and to test whether its WL+FR-induced increase depends on a shoot-borne HY5 source. With a Col-0 scion, providing shoot-borne HY5, HY5-GFP in lateral root primordia increased under shoot WL+FR, whereas with a *hy5hyh* scion this increase was absent (Figure 7). This indicates that the root-expressed HY5 response to WL+FR depends on shoot-expressed HY5.

**Figure 7:**
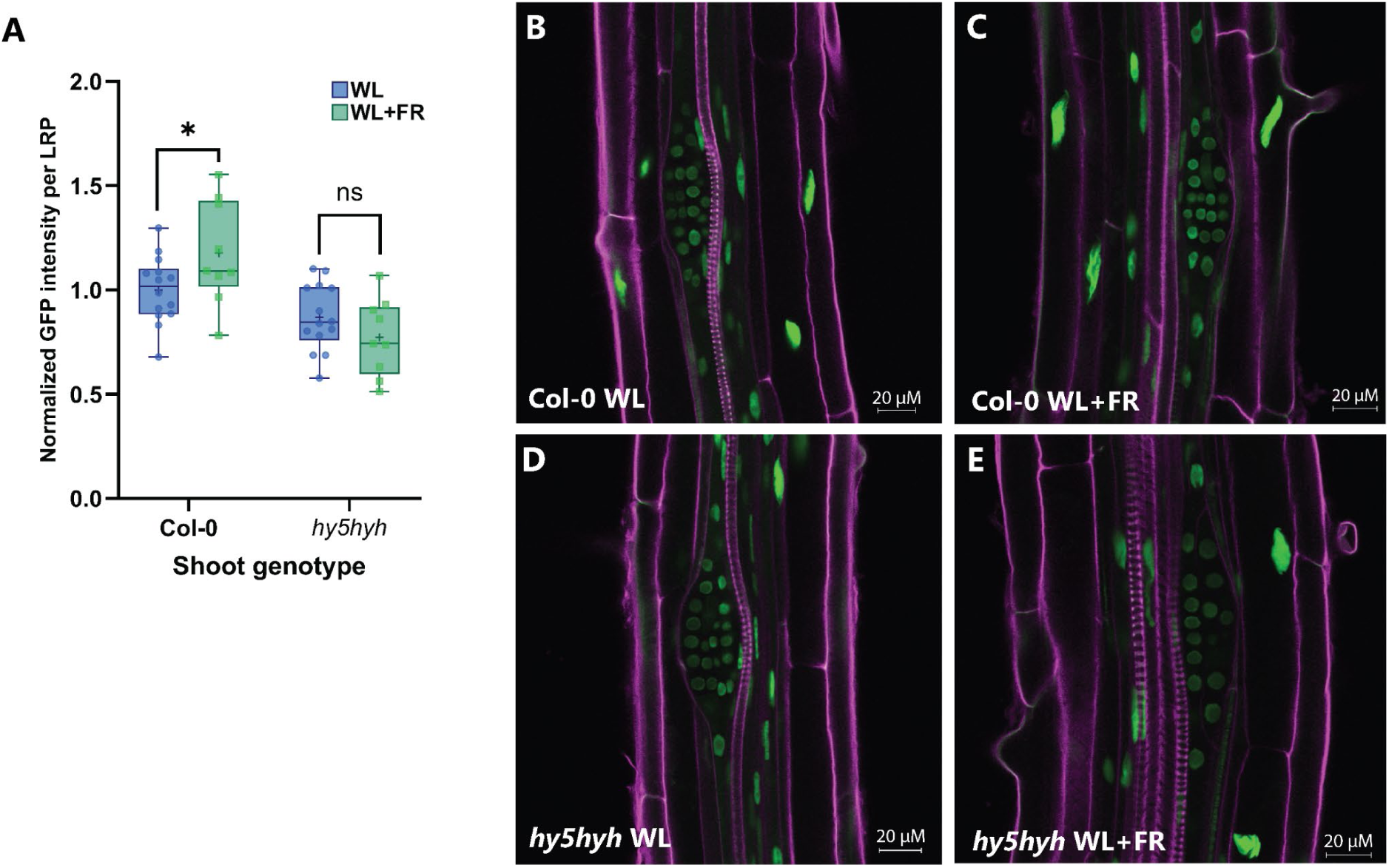
HY5-GFP intensity in LRP of grafted seedlings with Col-0 or *hy5hyh* scion grafted on a *hy5-2 pHY5:HY5-GFP rootstock.* (A) Normalized HY5-GFP intensity plotted per graft combination and treatment. Mean HY5-GFP intensity is normalized against Col-0 WL average. After grafting, seedlings were treated with WL or WL+FR from day 11 up to fixation after 14 days. Data is pooled from two subsequent experiments, N = 14 (Col-0 WL), N = 9 (Col-0 WL+FR), N = 14 (*hy5hyh* WL), N = 9 (*hy5hyh* WL+FR). Statistical differences are calculated between treatments per shoot genotype with a Student’s t-test (B) Representative confocal image of a single z-layer in the middle of the LRP of a Col-0 WL scion graft. GFP channel in green and cell wall staining in purple. (C) Representative confocal image of a single z-layer in the middle of the LRP of a Col-0 WL+FR scion graft. (D) Representative confocal image of a single z-layer in the middle of the LRP of a *hy5hyh* WL scion graft.(E) Representative confocal image of a single z-layer in the middle of the LRP of a *hy5hyh* WL+FR scion graft.

## Discussion

Aboveground competition for photosynthetically active light reduces lateral root development through a reduction in lateral root emergence that depends on HY5 (van Gelderen et al. 2018). HY5 has also been shown to move from shoot to root (Chen et al. 2016). Whether these two observations represent parts of the same pathway that explains light quality-driven plasticity of lateral root development was unclear. Here, we provide evidence that HY5 indeed stabilises in the shoot followed by translocation to the root that leads to local accumulation in the lateral root primordia. Using shoot-inducible *pCAB1:HY5* lines, we show that HY5 arrives in the root and unloads at the protophloem pericycle (Figures 1 & 2). HY5-GFP signal at the unloading zone was increased by shoot-perception of WL+FR (Figure 4), and the shoot-specific expression in *pCAB1:HY5* seedlings showed that shoot-borne HY5 is sufficient to rescue the lateral root density phenotype of *hy5-215* knockout roots (Figure 6). In addition, grafting *hy5hyh* scions onto *hy5-2 HY5-GFP* rootstocks abolishes the HY5-GFP accumulation in the lateral root primordia that was observed with Col-0 scions (Figure 7) and that were previously reported in intact *hy5-2 HY5-GFP* seedlings (van Gelderen et al. 2018). These data together support a model in which shoot-sensed WL+FR increases HY5 in the shoot, which translocates to the root, where shoot-derived HY5 is required for the increase of HY5 in lateral root primordia that regulates the reduction of lateral root emergence.

Several aspects of this model require closer examination, starting with the initial step: how shoot-perceived light conditions translate into HY5 delivery to the root. Shoot perception of WL+FR resulted in higher HY5 levels in shoot tissue (Figure 3), and in parallel we observed a higher intensity of HY5 transport events under WL+FR compared to WL (Figure 4, 5). This conditional stabilization of HY5 is most likely due to inhibition of the SPA/COP1 complex by phytochromes (Lu et al. 2015; Sheerin et al. 2015). Both *phyA* and *phyB* knockout mutants did not have a lateral root reduction in WL+FR, while showing opposing constitutive phenotypes (van Gelderen et al. 2018). Unlike phyB, phyA is activated by WL+FR, thus phyA is most likely the main actor in HY5 stabilization in this context. Despite this conditional stabilization under low R:FR, some shoot-induced HY5 was unloaded in the root even under WL conditions (Figure 4), with HY5-GFP protein detectable in root samples (Figure 5). Nevertheless, it is possible that the *pCAB1:HY5* construct in these experiments led to some overexpression of HY5 in shoot tissues. Indeed, shoot and root phenotyping of this *pCAB1:HY5* line indicates that it has a similar phenotype to a *35S:HY5* overexpression line published previously (van Gelderen et al. 2018). We propose that when HY5 levels increase, either through enhanced expression or through stabilisation in FR-enriched light, more of it will accumulate for translocation into the root system. Since phloem is the plant’s main route for translocation of solutes from shoot-to-root, we anticipate that this is also where HY5 is translocated. The localisation of HY5 signal (Figures 1-4) is consistent with this.

The phloem-loading mechanism of HY5 is not clear. Based on sparse literature, HY5 and HY5-GFP/YFP are small enough to move passively from phloem companion cells into the sieve elements (Balachandran et al. 1997). Although there could theoretically also be dedicated (active) transport of HY5, there are no studies to support that. Thus, protein phloem loading, whether active or passive, requires further study. In the primary root, unloading of HY5-GFP was restricted to the end of the protophloem, at the interface with the protophloem pericycle, where HY5 is released as the phloem stream ends (Fig. 1, Fig. S5 (Jelínková et al. 2026)). This region has plasmodesmata with an unusually high size exclusion limit (>120 kDa; (Ross-Elliott et al. 2017) and transport of HY5-GFP, as well as native HY5 would be compatible with passage through them. Furthermore, HY5 unloads with burst-like dynamics that are reminiscent of what has been described for general phloem unloading (Ross-Elliott et al. 2017; Jelínková et al. 2026), further supporting basic phloem-driven transport.

It is unclear how shoot-derived HY5 drives its own accumulation in the LRP once it has reached the roots. The observations that HY5 unloading may be restricted to the protophloem pericycle, and that shoot-derived HY5 does not seem to reach the LRP very well, raises the question of how shoot-derived HY5 influences accumulation in the LRP, thereby inhibiting lateral root development. The ability of shoot-derived HY5-YFP to complement the reduced lateral root density phenotype of the *hy5-215* mutant indicates that shoot-derived HY5 is active in roots and that its influence is not limited to the unloading zone. Our findings show that shoot-borne HY5 is necessary to drive HY5 accumulation in the LRP (Figure 7). This suggests that shoot-derived HY5 may promote local HY5 production or stabilisation in the root, perhaps more than supplying the LRP pool directly. HY5 has been shown to bind to its own promoter and activate its own expression, together with either CAM7 or PIF3, or in tomato, with BBX20 and BBX21 (Chen et al. 2013; Abbas et al. 2014; Yang et al. 2022). This regulation has the potential to amplify the HY5 signal in target tissues, such as the LRP.

Assuming that HY5 is indeed transported rootwards via the phloem and unloaded in the phloem unloading zone, presents a non-selective route that would not be unique to HY5. However, HY5 is a unique transcription factor, that can potentially bind the promoters of thousands of Arabidopsis genes (Lee et al. 2007), and at the same time requires cofactors (for example BBX proteins), to determine its specificity, since it lacks a traditional transactivation domain (Burko et al. 2020; Bursch et al. 2020). This makes HY5 a powerful signal that achieves specificity by being translated by cofactor transcription factors, thus allowing HY5 to act as an effective long-distance signal of light quality and neighbour shade.

## Methods

### Plant material and growth conditions

Existing mutant lines that were used in this study are: *hy5-215* (Oyama et al. 1997)*, hy5-2* (van Gelderen et al. 2018) *hy5-2 pHY5:HY5-GFP* (Zhang et al. 2017), *hy5-2 35S:HY5-GFP* (Zhang et al. 2017). For this study several new transgenic lines were constructed: *Col-0 pCAB1-YFP-HY5-HIS* and *hy5-215 pCAB1-YFP-HY5-HIS*. Col-0 *pCAB1-XVE:HY5-YFP* and *hy5-215 pCAB1-XVE:HY5-YFP*, *hy5-2 pHY5:HY5-YFP*. For the experiments in Figures 2 and 4, line Col-0 *pCAB1-XVE:HY5-YFP* #15-3 and *hy5-215 pCAB1-XVE:HY5-YFP* #2-4 were used. For the experiments in figure 5, A cross was used to generate with an original Col-0 *pCAB1-XVE:HY5-YFP* line (#14-2), with *hy5-215* to generate a *hy5-215 pCAB1-XVE:HY5-YFP* line with identical insert. In general, seeds were sterilized using a 1-2 hour treatment with chlorine gas. Seeds were sown on ½ MS 0.1% MES, pH 5.8, 0.8 % plant agar plates, (Greiner plates 12,5x12,5x1,75 cm) and then the plates were put in the dark at 4°C for six days for seed stratification. A D-root (Silva-Navas et al. 2015) insert was placed and the root part of the plate was covered by a black cover (see (van Gelderen et al. 2018)). *Arabidopsis thaliana* plants were grown in a controlled environment growth chamber with a 16-h-light/8-h-dark cycle, temperature of 20°C and a light level of PAR ∼140 µmol/m^2^/s. Supplemental far-red treatments were performed using Phillips GreenPower LED research modules (far red 730 nm, 24Vdc/10W) to achieve a R:FR ratio of 0.1 in combination with fluorescent tube lighting and 0.5 in combination with white LEDs. In total ∼60-70 µE of far-red light was supplied. Light measurements were performed with a UPRTEK PGN200 spectrophotometer. All phenotyping experiments were performed as described in (van Gelderen et al. 2018, 2025).

### Molecular cloning and construction of transgenic lines

For all primers used in this study, consult supplementary table 1. Tissue specific non-inducible lines were created by using the system described by (Michniewicz et al. 2015), using the *pCAB:YFP-Gateway* destination vector and recombining it with *pDONR207-HY5-HIS* (van Gelderen et al. 2025). Entry vectors, destination vectors and gateway cloning were all transformed into electrocompetent dH5a *E.coli*. We made a transgenic, estradiol-inducible, tissue-specific HY5 lines Col-0 *pCAB1-XVE:HY5-YFP* and *hy5-215 pCAB1-XVE:HY5-YFP*), by using the estradiol-inducible system of (Siligato et al. 2016), and a set of promoter cassettes from the group of Ari-Pekka Mahonen at Helsinki University. (https://www.helsinki.fi/en/researchgroups/plant-growth-dynamics/tools/tissue-specific-inducible-vectors). We constructed a *p1R4-pCAB1-XVE* vector by subcloning and ligating the *pCAB1* fragment from the aforementioned Michniewicz vectors. *p1R4-pCAB1-XVE* was then combined in a multisite gateway reaction with *pDONR207-HIS-HY5* (van Gelderen et al. 2025) *and p2R3-YFP-3AT* entry vectors in a *pGII-R3R4* entry vector (Siligato et al. 2016) (Addgene). *pHY5:HY5-YFP* vectors were constructed by subcloning a promoter region of 1500bp upstream of the start codon of *HY5*, into a pJET cloning vector. The existing *pDONR:HY5* clone was then used to sublime a new *HY5* fragment, and finally a *YFP-3AT* fragment was subcloned from the Siligato vector and all three fragments were ligated into a pGreenII destination vector with hygromycin resistance cassette. Destination vectors were then transformed to dh5a *E.coli*, and subsequently prepped and transformed into AGL1 psoup *A. tumefaciens.* For all transgenic lines, the transformation of *Arabidopsis* flowering plants was done via a published floral dip method (Davis et al. 2009). Transformed T1 seeds were selected with Norflurazone and propagated. ∼20 T2 lines were selected based on mendelian segregation to select for single inserts and also tested with epifluorescence microscopy for leaky expression and fast and correct induction. ∼3 T3 lines were selected for homozygosity with hygromycin and again tested for proper, fast non-leaky induction. Additionally, the location of the insert of line Col-0 *pCAB1-XVE:HY5-YFP* #14-2-4 was then determined by tailing PCR and subsequently crossed with *hy5-215* to yield line *hy5-215 pCAB1-XVE:HY5-YFP #14-2-4*.

### Grafting experiments

Seeds were sown 25 per row, two rows per plate and after 3d stratification were germinated in WL control conditions with the root part uncovered. After 5 days of growth, seedling hypocotyls (scions) of *hy5-2 35S:HY5-GFP* or *hy5-2 pHY5:HY5-GFP* (Figure 1) or, Col-0 or *hy5hyh* (Figure 7) were grafted onto Col-0 (Figure 1) or *hy5-2 pHY5:HY5-GFP* (Figure 7) rootstocks. Scions and rootstocks were cut with a sterile blade 1/3 above the shoot-root junction and the cut ends were joined in a 2 mm long, 0.3 mm diameter silicone tube (HelixMark, Freudenberg Medical). Grafts were assembled on Amersham^TM^ Hybond-N^+^ membrane strips placed on square petri dishes with 30 mL 1.5% agar, ½ MS and 1 g/L MES medium with 16 grafts per plate arranged on two membrane strips. Seedlings were then allowed to recover for 6 days in WL with the roots exposed to light (no dark-root cover). Successfully grafted seedlings were removed from the tubes, any adventitious roots were excised, and seedlings were transferred to new d-root plates containing 25 ML of ½ MS, 1% agar and 1 g/L MES medium, arranged in a single row of 14 seedlings with a d-root insert and root covered from the light. FR supplementation began on day 11, and after 14 days seedlings were fixed with 4 % PFA, cleared with ClearSee and stained with Calcofluor White for confocal imaging (Ursache et al. 2018). Confocal imaging was performed using a Leica Stellaris 5 confocal microscope (Leica Microsystems) using a 40x objective acquiring tile scans (Figure 1) and z-stacks (Figure 7). GFP fluorescence was excited at 485 nm with a white light laser and detected between 490-839 nm. Calcofluor white cell wall staining was excited at 405 nm and detected between 426-482 nm. The z-layers making up the middle of the LRP were combined in a maximum projection and analyzed with ICY. Nuclei in the LRP were semi-automatically detected with the HK-means plugin, and manually corrected with the area ROI tool if needed. Mean intensity was normalized against the mean of the Col-0 scion grafts in control WL.

### Western Blotting

Seedlings were grown as described above. At five days post-germination, seedlings were either maintained under WL conditions or transferred to WL supplemented with FR (WL+FR). On day 6, HY5-YFP expression was induced by applying 5 µM β-estradiol in 0.1% agarose to the shoot, while an equivalent amount of DMSO in 0.1% was used for mock treatment. To inhibit protein translation locally, cycloheximide (CHX; 50 or 100 µM) was applied separately to either the shoot or root. Shoot and root tissues were harvested separately 24 h after treatment. Approximately 10 shoots and 20 roots per sample were frozen in liquid nitrogen and ground with beads for 1 minute, re-cooled in liquid nitrogen, and then ground for an additional 1 minute. Proteins were extracted in 2× Laemmli buffer (4% SDS, 120 mM Tris-HCl pH 6.8, 10% glycerol, 0.05% bromophenol blue, and 5% β-mercaptoethanol) supplemented with protease inhibitor and 10 µM bortezomib. Shoot samples were extracted in 200 µL extraction buffer and root samples in 100 µL. Samples were heated at 95°C for 5 minutes and centrifuged for 1 minute at maximum speed. Proteins were separated by SDS-PAGE using 12% resolving gels and 4.8% stacking gels (4x buffer, 0,4% SDS, 1,5 M Tris-HCl pH 8,8). For each sample, 6 µL of shoot protein extract and 14 ul of root protein extract were loaded onto the gel. Electrophoresis was initially performed at 60 V until the samples entered the resolving gel, then continued at 130 V, using standard Tris-SDS-Glycine-SDS buffer (1% SDS, 0,25M Tris-glycine, pH 8,6). Proteins were transferred to the membrane by wet transfer for 60 minutes at 100 V, using standard Tris-Glycine-SDS buffer with 20% ethanol. Membranes were blocked overnight at 4°C in 1× TBS containing 5% (w/v) milk powder. YFP-tagged proteins were detected using a custom polyclonal anti-GFP primary antibody (Roth et al. 2018)(1:2,500 for roots and 1:10,000 for shoots) diluted in 1× TBS containing 0.5% (w/v) milk powder. Membranes were subsequently washed four times for 10 minutes with 1× TBS containing 0.1% Tween. Then, they were incubated for 1 h at room temperature with an HRP-conjugated anti-IgG secondary antibody (Agrisera AS09 602) in 1× TBS containing 0.5% (w/v) milk powder. The secondary antibody was used at a dilution of 1:5000 for root samples and 1:20,000 for shoot samples. Membranes were then washed four additional times for 10 min with 1× TBS containing 0.1% Tween. Signals were detected using ECL (Cytiva Amersham ECL Prime), and membranes were subsequently stained with 0.1% Ponceau S in 5% acetic acid as a loading control.

### Live confocal microscopy imaging

Seeds were sown and grown according to the protocol described above. Seedlings growing with supplemental far-red were exposed to treatment 1 day post germination. At 5 days post germination, a 0.1% agarose droplet containing 5 µM β-estradiol (dissolved from a 5 mM stock in DMSO) was carefully pipetted on the seedling shoot in the morning, ∼ 4 hrs post-dawn. Seedlings were then placed back into the growth chamber for 6 hours. Induced seedlings were then carefully placed on an Ibidi chambered coverslip (https://ibidi.com/chambered-coverslips/314--slide-1-well.html) together with a small droplet of ½ MS liquid medium. On top of the hypocotyl and root parts, an agar block cut from the ½ MS plate was placed, and the chamber was closed with the lid and Leucopore tape. Seedlings were imaged in Zeiss LSM880 confocal microscope, customized by CARL ZEISS Česká republika, to be positioned at 90 degrees to allow imaging according to the principles of (von Wangenheim et al. 2017). A ring of blue-red LED lights was placed around the microscopy table to supply growth light during the day part of the experiment. 4-5 seedlings root tips were imaged at the same time overnight (∼16-20 hrs) using a 20x air objective, 514 laser, and 520-560 bandpass filter, in tilescan mode. Z-stacks were then processed by ZEN Blue and ICY (De Chaumont et al. 2012). Max-Z projected images were then quantified by ICY, using the area ROI tool.

### Sample fixation and confocal microscopy

Seedlings used in Figure 3 were fixed using 4% formaldehyde, washed twice with PBS buffer, cleared using clearsee, and stained according to (Ursache et al. 2018). Samples for Figure S5 were also fixed with 4% formaldehyde, but subsequently cleared with clearsee alpha (sodium sulfite) (Kurihara et al. 2021). Figure 3 confocal microscopy was performed on a Zeiss LSM7, using a 514 laser, and 525-560 bandpass filter with a 40x oil objective. Figure S5 confocal microscopy was performed with a Zeiss LSM 990 in normal confocal mode, using a 514 laser and 525-560 bandpass filter and 40x oil objective.

## Supplemental figures

**Figure S1:**
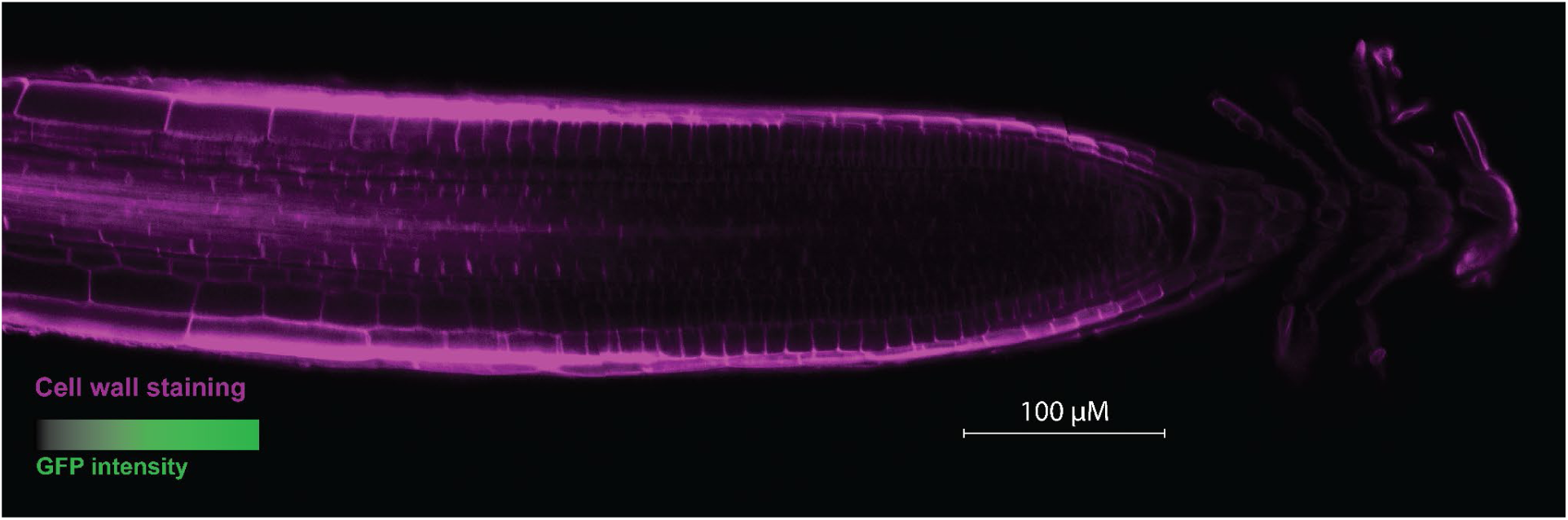
Root tip of homograft of Col-0/Col-0. Cell-wall staining in purple and GFP intensity in green, same settings as for grafts in Figure 7 (not visible in Col-0/Col-0 graft). Scale bar is 100 μM.

**Figure S2:**
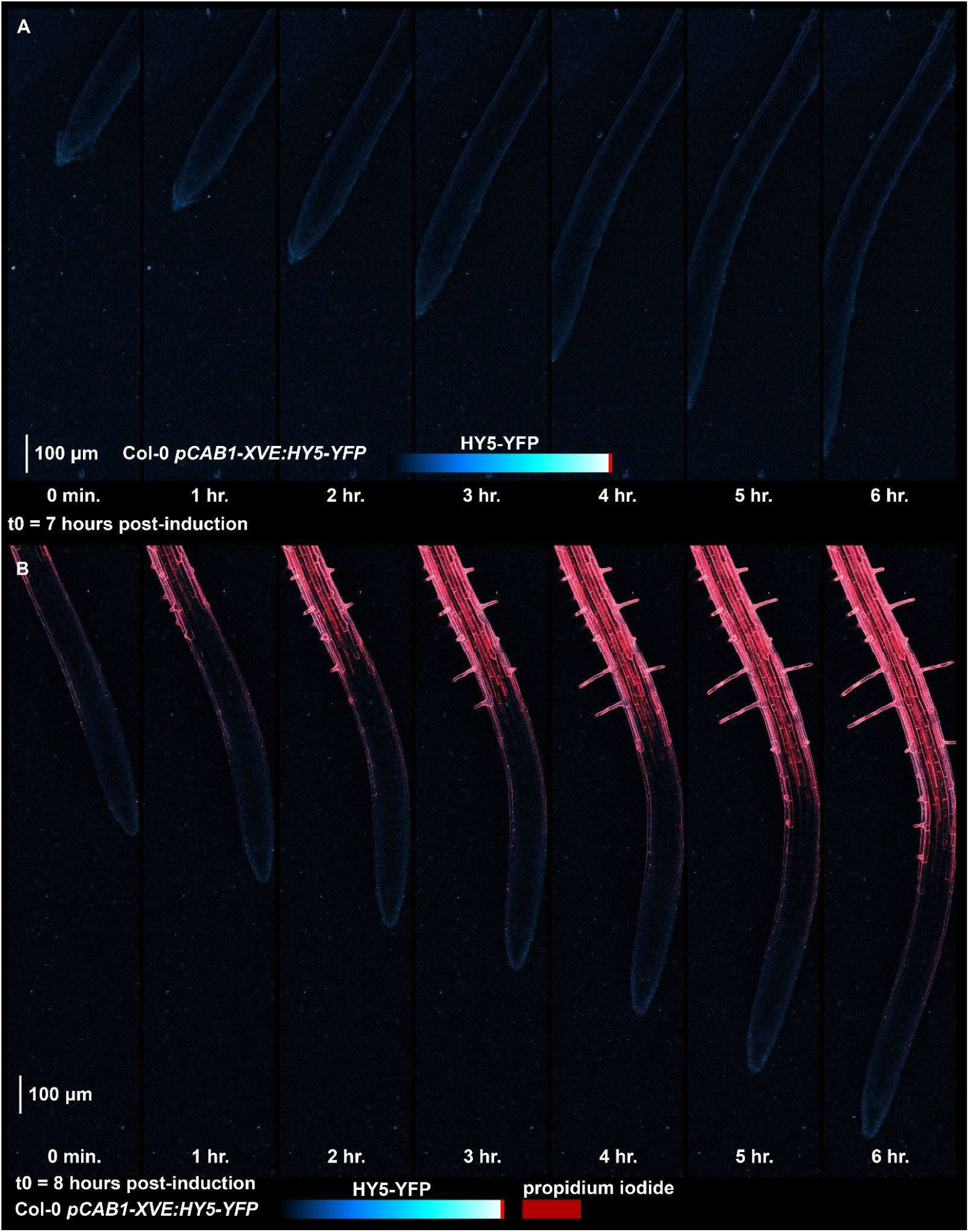
Mock DMSO treatment on shoot or direct beta-estradiol treatment on root did result in HY5 shoot-root transport. Confocal images of Col-0 *pCAB1-XVE:HY5-YFP* lines treated not with beta-estradiol, but with DMSO (A), or treated with estradiol directly on the root (B).

**Figure S3:**
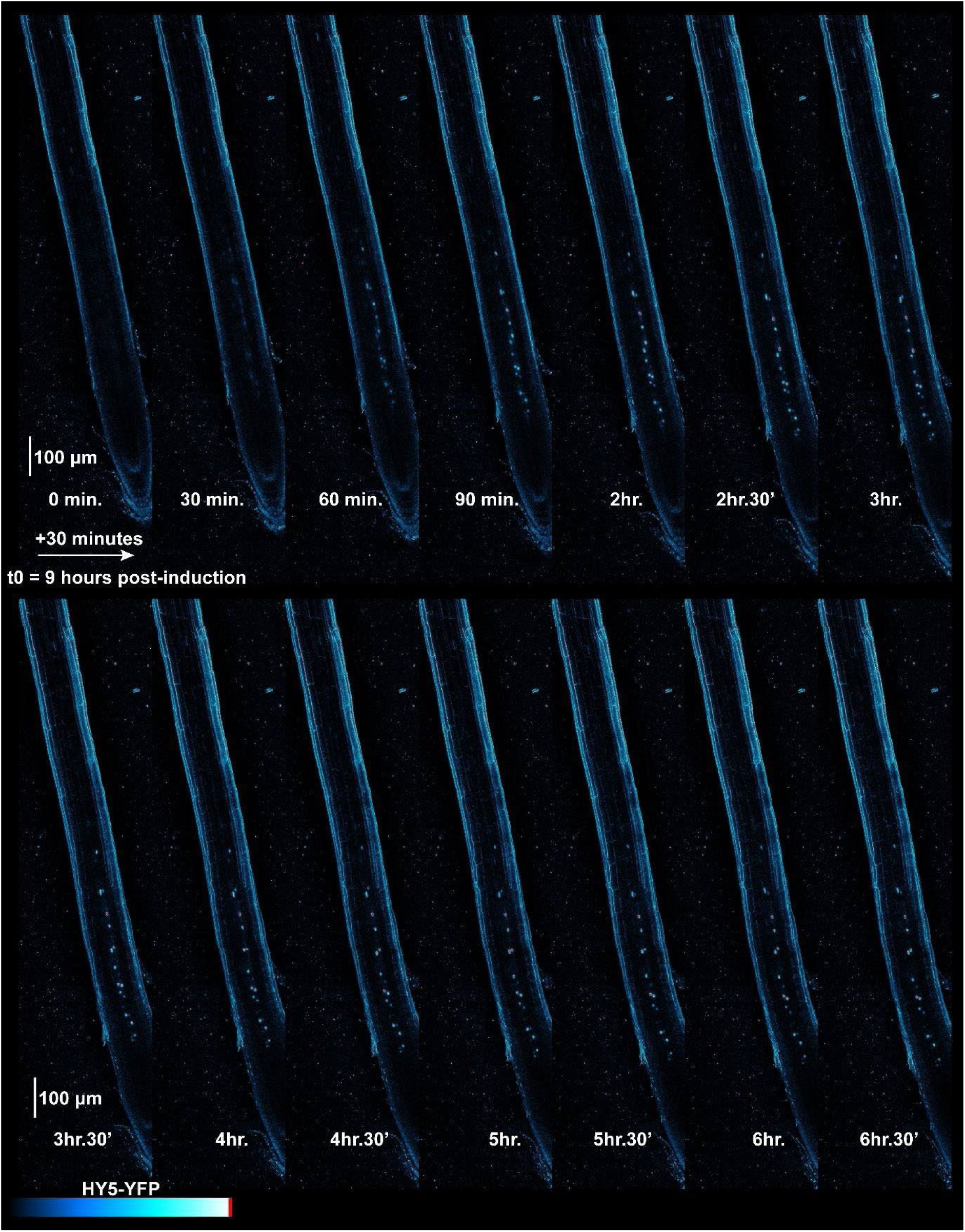
HY5 Transport of *hy5-215 pCAB1-XVE:HY5-YFP.* Confocal image series of experiment set up in the same manner as Figure 1, with the difference being the genetic background of the inducible construct.

**Figure S4:**
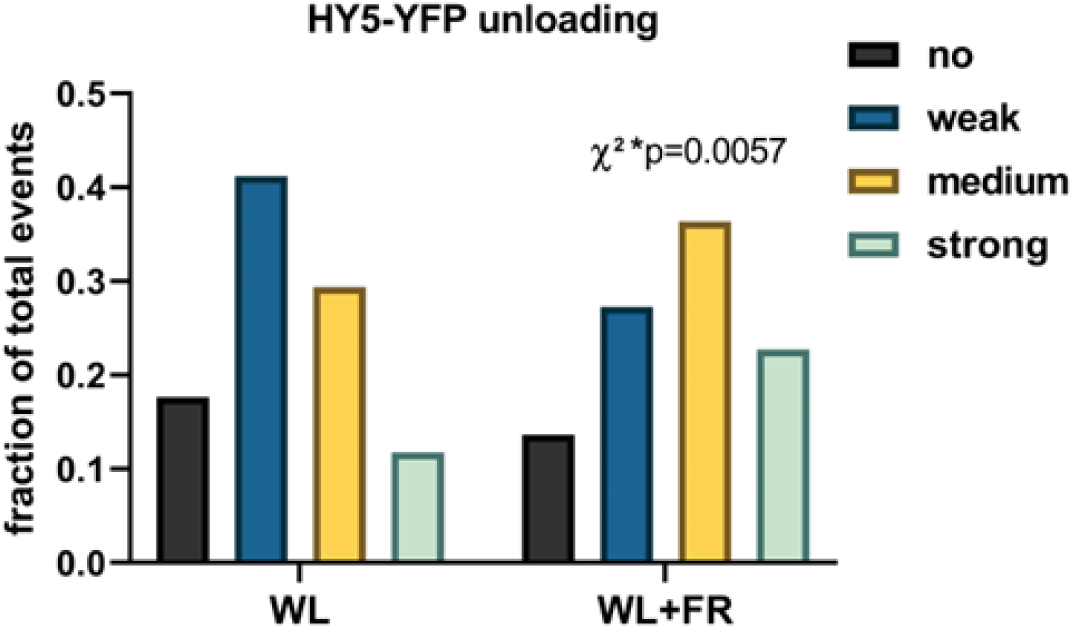
More HY5 transport events occur in supplemental far-red treatment. Manual counting of HY5 transport events frequency, of the images in the dataset of Figure 4. Signals were assessed on their relative strength and scored in one of the categories (no, weak, medium or strong transport signal). A chi-square test compared the ratios of the WL+FR to the WL and it was found to be significantly different (p=0.0057; n (WL) = 36, n (WL+FR) = 25).

**Figure S5:**
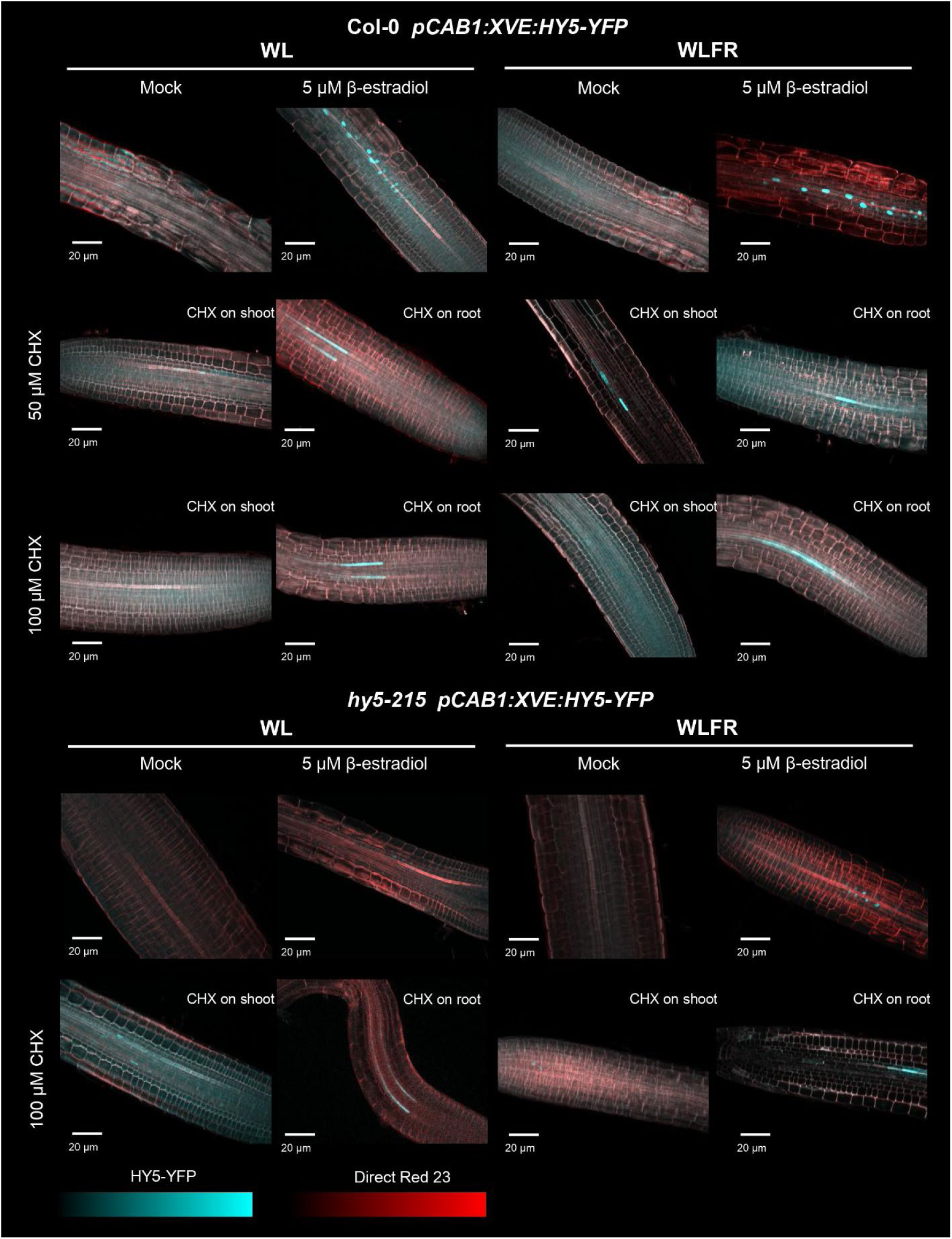
HY5 transport signal is not dependent on inhibition protein of translation in the root. Confocal microscopy Images of root tips of seedlings treated in the same way as in Figure 5. Col-0 *pCAB1-XVE:HY5-YFP* and *hy5-215 pCAB1-XVE:HY5-YFP* lines were induced with 5 µM beta-estradiol on the shoot, and additionally the translation inhibitor cycloheximide (CHX) was applied specifically on the shoot or root. After 24 hours, seedlings treated with either WL or WL+FR were fixed in formaldehyde and cleared via Clearsee. Shown are representative images, the number of biological replicates is between 10 and 20.

## Supplementary Movies

Movie S01: Complementing Figure 2 *Col-0-pCAB1-XVE-HY5-YFP*

Movie S02: Complementing Figure 2 *hy5-215-pCAB1-XVE-HY 5-YFP*

Movie S03: Complementing Figure 4 Col-0 pCAB1-XVE-HY5-YFP_mock

Movie S04: Complementing Figure 4 Col-0 pCAB1-XVE-HY5-YFP_WL

Movie S05: Complementing Figure 4 Col-0 pCAB1-XVE-HY5-YFP_WL+FR

